# Enjoy The Violence: Is appreciation for extreme music the result of cognitive control over the threat response system?

**DOI:** 10.1101/510008

**Authors:** Rosalie Ollivier, Louise Goupil, Marco Liuni, Jean-Julien Aucouturier

**Affiliations:** STMS UMR9912 (IRCAM/CNRS/Sorbonne Universit), Paris (France)

**Keywords:** Music, Emotion, Metal, Fear, Threat, Cognitive load

## Abstract

Traditional neurobiological theories of musical emotions explain well why extreme music such as punk, hardcore or metal, whose vocal and instrumental characteristics share much similarity with acoustic threat signals, should evoke unpleasant feelings for a large proportion of listeners. Why it doesn’t for metal music fans, however, remains a theoretical challenge: metal fans may differ from non-fans in how they process acoustic threat signals at the sub-cortical level, showing deactivated or reconditioned responses that differ from controls. Alternatively, it is also possible that appreciation for metal depends on the inhibition by cortical circuits of a normal low-order response to auditory threat. In a series of three experiments, we show here that, at a sensory level, metal fans actually react equally negatively, equally fast and even more accurately to cues of auditory threat in vocal and instrumental contexts than non-fans. Conversely, cognitive load somewhat appears to reduce fans’ appreciation of metal to the level reported by non-fans. Taken together, these results are not compatible with the idea that extreme music lovers do so because of a different low-level response to threat, but rather, highlight a critical contribution of higher-order cognition to the aesthetic experience. These results are discussed in the light of recent higher-order theories of emotional consciousness, which we argue should be generalized to the emotional experience of music across musical genres.

## Introduction

Our capacity to perceive affects in music has been the subject of impassioned psychology and neuroscience research in the past two decades (Blood and Zatorre 2001; Juslin and Västfjäll 2008). While music was once believed to have a “language of emotions” of its own, separate from our species’ other expressive capacities (McAlpin 1925), today’s dominant view of musical expression construes it as in many ways continuous with natural languages (Patel 2007). Musical emotions are studied as communicative signals that are encoded in sound by a performer, then decoded by the listening audience (Juslin and Laukka 2003), for whom hearing music as expressive involves registering its resemblance with the bodily or vocal expressions of such or such mental state (Juslin and Västfjäll 2008). For instance, joyful music is often associated with fast pace and animated pitch contours (as is happy speech), melancholic music with slower and flatter melodic lines and dark timbres (as is sad speech), and exciting music with high intensity and high levels of distortion and roughness (as may be an angry shout) (Juslin and Laukka 2003; Ilie and Thompson 2006; Escoffier et al. 2013; Blumstein et al. 2012).

Seeing musical expression as a culturally-evolved phenomena based on a biologically-evolved signaling system (Bryant 2013) allows to explain much of people’s typical affective responses to music. Just like vocalizations, music which signals happiness or affiliation may be appraised positively or lead to positive contagion (Miu and Balteş 2012); sad music may be reacted to with empathy, and make people sad or moved (Vuoskoski and Eerola 2017). Similarly, humans, and many non-human animals, produce harsh, rough and nonlinear sounds when alarmed (Anikin et al. 2018). In ecological situations, such sounds trigger stereotypical fear and avoidance behaviors (e.g., in conditioning paradigms - Den et al. 2015), are strong prioritized in sensory processing (Asutay and Västfjäll 2017), and evoke activity in areas linked to the brain’s threat response system (Arnal et al. 2015). It is therefore no surprise that “extreme” music such as punk, hardcore or metal, whose vocal and instrumental characteristics share much acoustic similarity with threat signals, should evoke for a large proportion of listeners feelings of anger, tension and fear (Rea et al. 2010; Blumstein et al. 2012; Thompson et al. 2018), impair their capacity to cope with simultaneous external stress (Labbé et al. 2007) and trigger reactions of avoidance and a desire to stop listening (Bryson 1996; Thompson et al. 2018). Decoding extreme music as an auditory signal of danger or threat, these listeners (as one respondent quoted in Thompson et al. 2018) literally *“[…] cannot understand how anyone finds this music pleasant to listen to”*.

Some listeners obviously do, though. Extreme music, and most notably metal music, is a thriving global market and subculture, with strongly engaged communities of fans (Guibert and Guibert 2016). Despite long-lived stereotypes that listeners who engage with metal music do so because of a psycho-socially dysfunctional attitude to violence and aggression (Stack et al. 1994; Bodner and Bensimon 2015; Sun et al. 2017), it is now well-established that listeners with high-preference for metal music do not revel in the strongly negative feelings this music usually induces in non-metal fans. Rather, metal music fans report that the music leads them to experience a wide range of positive emotions including joy, power and peace (Thompson et al. 2018) and no increase of subjective anger (Gowensmith and Bloom 1997). In fact, following an anger-induction paradim, Sharman and Dingle (2015) report that listening to 10 minutes of violent metal music *relaxed* metal music fans just as effectively as sitting in silence. It therefore appears that metal music fans do not process the threat-signaling features of violent music to the same outcome as non-metal fans. It is not that they enjoy the threat; rather, they do not experience threat at all.

While traditional, neurobiological views of emotions link the emergence of emotional feelings, such as that of experiencing fear, to the operation of innately-programmed, primarily sub-cortical brain systems, such as those centered on the amygdala (Panksepp 2004), more recent cognitive frameworks tend to separate the activation of such circuits from that of higher-order cortical networks that use inputs from sub-cortical circuits to assemble the emotional experience (LeDoux and Pine 2016; LeDoux and Brown 2017). In short, while first-order threat responses may contribute to higher-order feeling of fear, they do not unequivocally constitute it : on the one hand, defensive survival circuits may be activated by subliminally-presented threatening visual stimuli and generate behavioural or autonomic threat response patterns even in the absence of subjective fear (Vuilleumier et al. 2001; Whalen et al. 2004; Diano et al. 2017); on the other hand, bilateral damage to the amygdala may interfere with bodily responses to threats, while preserving the conscious experience of fear (Feinstein et al. 2013; for a discussion, see Fanselow and Pennington 2018). In sum, autonomic, behavioural and primitive responses to threat stimuli appear to be neither necessary nor sufficient for the conscious experience of fear to emerge.

The existence of two populations, metal fans and non-fans, that respond to identical cues of auditory threat with radically different emotional experience (pleasure/approach, or fear/avoidance) provides a compelling ecological situation in which to study how first-order and high-order processes interact to create emotional states of consciousness. On the one hand, it is possible that metal fans differ from non-fans in how they process threat signals at the first-order/sub-cortical level. Just like clinical populations with specific phobias or social anxiety show increased amygdala reactivity to their trigger stimuli (e.g. pictures of spiders or fearful faces) even when presented outside of conscious awareness (McCrory et al. 2013; Siegel et al. 2017), metal fans may show deactivated responses to the cues of auditory threat constitutive of that musical genre, possibly as the result of positive conditioning (see e.g. Blair and Shimp 1992). If present, such first-order, bottom-up differences between fans and non-fans would not only predict a different late-stage read-out of the activity of the threat circuit (i.e. experiencing fear or not), but also different autonomic and behavioural responses to auditory roughness even beyond the realm of music (e.g. fans not reacting to angry voices as fast/as negatively as non-fans). On the other hand, it is also possible that the fans’ appreciation for metal music reflects a higher-order inhibition by cortical circuits of an otherwise normal, low-order response to auditory threat. In support of this dissociation, Gowensmith and Bloom (1997) found that, while metal fans listening to metal music reported feeling less angry than non-fans, both fans and non-fans reported similar levels of physiological arousal in response to metal music, suggesting that lower-order circuits reacted similarly in both groups. Conversely, a number of studies have shown that loading executive functions with visual attention (Pessoa et al. 2002), working memory (Van Dillen et al. 2009) or demanding arithmetic tasks (Erk et al. 2007) can lessen both the subjective evaluation, and amygdala response to negative stimuli. If they are involved in musical aesthetic experiences, we should predict that such higher-order, top-down processes would be more engaged for metal fans than non-fans during the emotional experience of metal music, and that loading these executive functions with a dual-task paradigm would lead to a failed inhibition of avoidance-related processes arising from the threat circuit, thereby lessening metal fans’ appreciation to the level experienced by non-fans.

In this article, we report on three experiments that aim to separate these two alternatives and to clarify the contribution of low- and higher-order processes in the emotional experience of metal music by fans and non-fans. We screened a total of 332 participants to constitute an experimental group of metal music fans that ranked low on appreciation for a control music genre (pop music) and a control group that ranked high on pop music but low on metal. To test the possibility of different low-order behavioural responses to threat cues, both groups rated the valence of vocal and musical stimuli presented with and without cues of arousal/roughness (**Experiment 1**). They were also subjected to a speeded spatial localization task with the same stimuli presented at different dichotic interaural time differences (ITDs) (**Experiment 2**). To test the contribution of higher-order inhibition to fans’ appreciation, we subjected both groups to a dual-task paradigm in which participants listened and rated their preference for both metal and pop music extracts while engaging in a demanding visual search task (**Experiment 3**). Our hypotheses, which we preregistered along with a basic data analysis strategy (Supplemental information SI2), were that groups would differ in Experiments 1 and 2 if metal appreciation is the result of different low-level processes, and would differ in Experiment 3 if it is the result of higher-order cognitive control over low-level processes.

## Experiment 1: valence rating task

A wealth of behavioural data suggests that cues of auditory threats, such as distortion, roughness and other non-linearities, are generally rated explicitly as low on valence. For instance, Arnal et al. (2015) finds that human listeners judge vocal, instrumental and alarm sounds re-synthesized to include temporal modulations in the [30,150] Hz range elicited more negative ratings, as well as faster response times, than similar unmodulated sounds; Blumstein et al. (2012) finds that musical soundtracks manipulated to include distortion were judged more negative and more arousing than control soundtracks. In the animal kingdom, marmots spend less time foraging after hearing alarm calls manipulated to include white noise than after normal or control calls (Blumstein and Recapet 2009). Here, we therefore take participants’ explicit ratings of the valence of short vocal and instrumental sounds (manipulated to induce roughness or not) as an index of affective responses to auditory threat in a generic, non-musical context, and test the hypothesis that such responses may be deactivated in metal fans.

### Methods

#### Participants

A total of 332 participants with normal self-reported vision and hearing were screened via an online questionnaire for their orientation toward a variety of musical genres, including metal, as well as a number of demographic variables. For each genre, participants had to indicate how much they enjoyed listening to such music, using a 7-point Likert scale. In addition, for genres rated above 5, they had to cite three of their favorite tunes for that genre. Genres listed in the survey were inspired by typical taxonomies of internet music services like Spotify (Pachet and Cazaly 2000), and included *blues, contemporary music, classical, French variety, electro, folk, jazz, metal, pop, rap/hip-hop, religious music, rock, soul/funk and world music*. Pop music was selected as a control genre for being not typically associated with strong cues of auditory threat, and for having high negative correlation with preference for metal music across the group (Pearson’s *r*=−0.12, n=332; FIgure 1).

**Figure 1.**
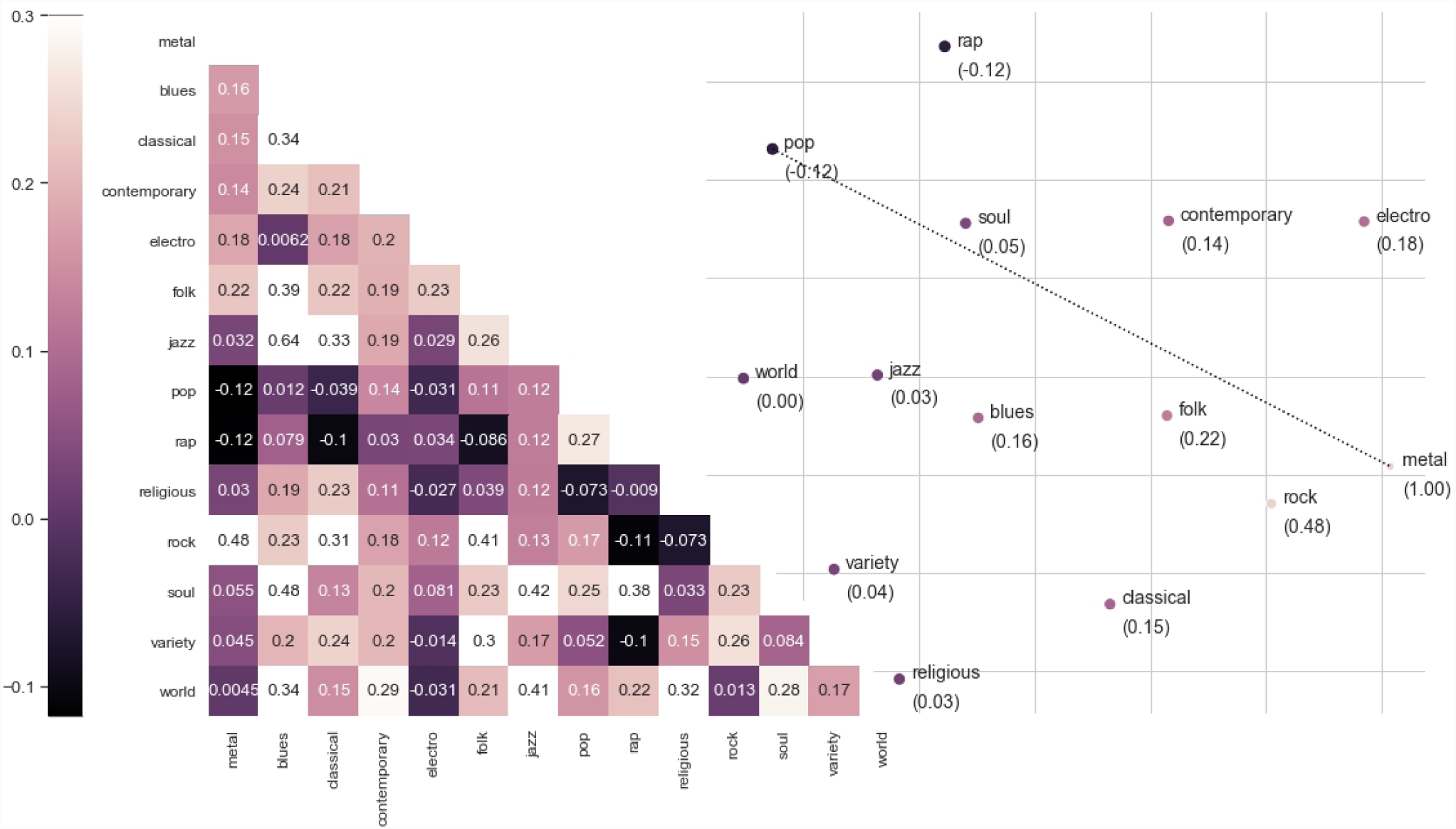
Relations between liking for musical genres in the N=332 participants screened for the study. Left: Correlation matrix between genres, labeled with Pearson’s *r* coefficients. Right: two-dimensional multidimensional scaling solution for the same correlations, each genre labeled with Pearson’s *r* correlation to metal. Participants who liked, or disliked, metal music tended to have similar attitudes to rock (*r* =0.48), and opposite attitudes to pop (*r* =−0.12) and rap (*r* =−0.12).

We then selected N=40 participants from the original pool, based on their orientation towards metal and pop music. 20 participants (male: 12; M=21.3yo,SD=2.7) who gave rates ≥ 6 for metal music and ≤ 4 for pop music were selected for the metal group, and 20 participants (male: 10; M=22.3yo,SD=3.2) who gave rates < 2 for metal and > 6 for pop music were selected for the control group. Metal fans did not statistically differ from controls in terms of age (mean difference M=−1.0y, 95% CI [-2.96,0.86], *t*(38)=−1.1, *p*=.27), musical expertise (mean practice difference M=4.9y, 95% CI [-1.7, 11.5], *t*(11)=1.63, *p*=.13) and musical engagement (mean listening difference M=−3.35hr/week, 95% CI [- 12.1,5.4], *t*(38)=−0.77, *p*=.44). 6 participants were eventually not able to participate in the study after they were included, leaving 17 participants in each group for the final sample (N=34).

#### Stimuli

Stimuli for the experiment consisted of 24 short, 1-second recordings of human vocalizations (12 original, 12 rough) and musical instruments (12 original, 12 rough). Original vocalizations were recorded by one female and two male actors instructed to shout/sing phonemes [a] and [i] at three different pitches (in the range [450,480], [570,600] and [520,570] Hz for females; [200,215], [250,270] and [315,340] Hz for males), with a clear, loud voice. Original musical instrument samples were extracted from the *McGill University Master Samples* sound library (MUMS; Opolko and Wapnick 1989), and included single note recordings of three wind (bugle, clarinet, trombone) and one string (violin) instrument, each performed at three different pitches. Both types of sounds were then manipulated with a digital audio transformation aimed to simulate cues of vocal arousal/roughness (ANGUS; Gentilucci et al. 2018; freely available: http://forumnet.ircam.fr/product/angus/). ANGUS transforms sound recordings by adding sub-harmonics to the original signal using a combination of f0-driven amplitude modulations and time-domain filtering, an approach known to confer a growl-like, aggressive quality to any vocal or harmonic sound (Tsai et al. 2010). Here, we used ANGUS to add 3 amplitude modulators at f0/2, f0/3 and f0/4 sub-multiples of the original sounds’ fundamental frequency (f0), and thus generated transformed “rough” versions of each of the 12 vocal and instrument original sounds, resulting in 24 vocal and 24 musical stimuli.

#### Procedure

Participants were presented with one block of 24 vocal and one block of 24 musical stimuli (counterbalanced), played through Beyerdynamics DT770 headphones. At each trial, participants were instructed to rate the perceived valence/approachability of the stimulus, using a 7-point Likert scale ranging from 1 (‘very negative’) to 7 (‘very positive’). Stimuli were presented in random order within each block, with an inter-stimulus interval randomized between 0.8-1.2 sec.

#### Preregistered analysis strategy

Participant ratings were analyzed with a rmANOVA, with participant group (metal/not) as a between participant factor and stimulus roughness (original/rough) as a within-participant factor.

### Results

There was a main effect of stimulus roughness on perceived approachability, with ANGUS-manipulated sounds judged more negative than original sounds (Figure 2; mean valence difference M=−0.41, 95% CI [−0.51, −0.33], *F*(1,32)=43.7, *p*=1.8.10^*-*^7, ges = 0.11). However, this effect of roughness did not interact with participant group: both metal fans and non-fans judged rough sounds less approachable than original sounds (mean valence difference M=−0.08, 95% CI [−0.26 0.10], *F*(1,32)=0.39, *p*=.53, ges = 0.001).

**Figure 2.**
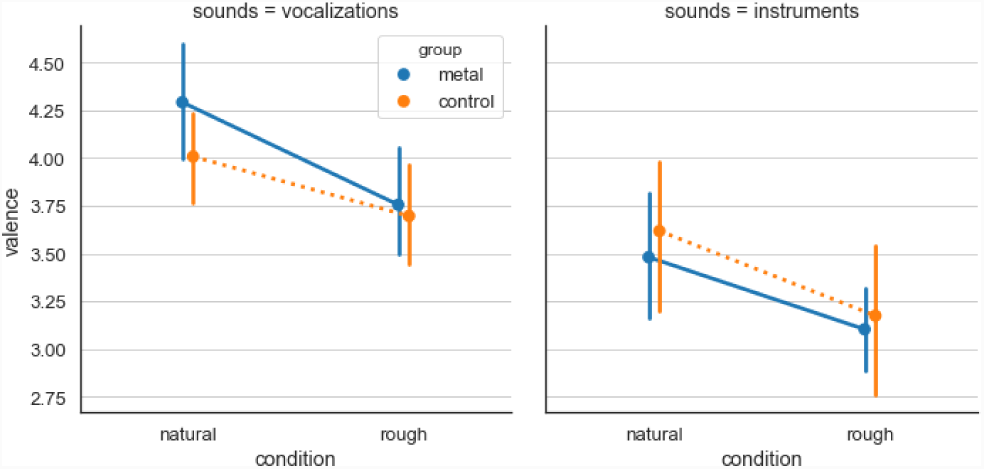
Effect of stimulus roughness on valence ratings (Exp.1), left: human vocalizations, right: musical instruments. Rough sounds were judged more negatively than original sounds, and metal fans did not report less negativity than non-metal fans for either rough vocal or musical sounds.

As an additional non-registered analysis, we also examined the effect of sound category (vocalization or instrument) on valence ratings: there was a main effect of category on valence ratings, with vocalizations judged more positive than musical instruments across conditions (Figure 2; mean difference M=0.59, 95% CI [0.42, 0.76],*F*(1,32)=26.67, *p*=1.2.10^−^5, ges = 0.21). However, this effect did not interact with either stimulus roughness (*F*(1,32)=0.023, *p*=.88), or participant group (*F*(1,32)=1.43, *p*=.24).

### Discussion

Our data replicates the finding that cues of auditory threat, as simulated here by amplitude modulations and the ANGUS software tool, are appraised as low on approachability / valence (Blumstein et al. 2012; Arnal et al. 2015). Interestingly, despite being grounded in biological signaling and the physiology of the vocal apparatus (Fitch et al. 2002), cues of auditory roughness elicited similar emotional evaluation regardless of whether they were applied to vocal or musical sounds, confirming that biological signaling indeed underlie part of the emotional reactions to musical sounds (Blumstein et al. 2012).

Critically for our hypothesis, however, metal fans did not report less negativity than non-metal fans for either rough vocal or musical sounds. Outside of a extreme musical context, metal fans therefore do not find rough sounds particularly pleasing and approachable, even with isolated instrument sounds. This does not support the idea that metal lovers do so because of altered or reconditioned affective responses to cues of auditory threat, but rather suggests that, outside of the culturally circumscribed musical context of metal music, such responses lead to the same behavioural outcome as in non-fans. Yet, because our rating task specifically targeted explicit affective judgments, it remains a possibility that low-level perceptual responses still differ in metal fans, but that these participants somehow compensate at the explicit level by relying on declarative knowledge, e.g. an awareness of the fact that rough sounds generally convey negative attitudes (e.g., shouts are often used in situations where people are angry). Thus, we ran a second experiment, to examine a purely perceptual process, sound localization, that although it is impacted by it, does not necessarily involve an affective evaluation of the stimuli, and operates on very short time scales that allegedly tap into more implicit mechanisms.

## Experiment 2: spatial localization task

Beyond the explicit negative appraisal of the stimuli, the rapid and accurate localization of danger is one of the main behavioural outcomes of the threat response system (Panksepp 2004). In previous work, Asutay and Västfjäll (2017) submitted participants to a visual search task and found that search times for low-salient targets decreased when these were preceded with task-irrelevant arousing sounds (dog growls and fire alarm). Similarly, Arnal et al. (2015) measured the speed and accuracy to detect whether normal vocalizations and screams were presented on participants’ left or right sides using ITD cues, and found participants were both more accurate and faster at localizing screams. Here, we implement a similar spatial localization experiment and use location speed and accuracy as an implicit index of threat responses in metal and non-metal fans, testing whether such behavioural outcomes are hypo-activated in metal fans.

### Methods

#### Participants

Experiment 2 included the same N=34 participants (metal: 17: non: 17) as Experiment 1.

#### Stimuli

Experiment 2 used the same 48 stimuli (24 voice, 24 instrument samples) as Experiment 1, with the same acoustic manipulation of roughness (ANGUS; Gentilucci et al. 2018) for half of the stimuli.

#### Procedure

We used a similar procedure as Arnal et al. (2015). Participants were presented with 15 repetitions of each stimuli (a total of 15×48=720 trials), played dichotically through Beyerdynamics DT770 headphones with an interaural time difference (ITD) indicative of either a left-field or right-field presentation. Prior to testing, stimulus ITD was individually calibrated for each participant using an 2-up-1-down staircase procedure, with a dichotically presented 300ms pure tone at fundamental frequency 700Hz. The initial ITD was 25samples (567.5*µs* at SR=44,1kHz), and the initial step size was 2 samples (45.4*µs*). This step size was halved (1 sample, 22.7*µs*) after the first inversion. Throughout the adaptive procedure, ITD values were constrained to a minimum of 22.7*µs* and a maximum of 567.5*µs*, and SOA was randomized between 0.8-1.2s. The procedure stopped after 12 inversions, and the final ITD was computed as the average ITD over the last 5 steps.

Testing then consisted of two blocks of 360 vocal and musical trials (counterbalanced, randomized with each block), dichotically presented at each participant’s fixed ITD, with a balanced, pseudo-random sequence of 360 left- and 360 right-field presentations. SOA was randomized between 1.4-1.9s. At each trial, participants were instructed to report their perceived field of presentation (left/right) as quickly as possible.

#### Preregistered analysis strategy

Similar to Arnal et al. (2015), we measured individual localization performance (d-prime), reaction times (RTs) and calculated a composite measure of efficiency, corresponding to the additive effect of individual z-score-normalized performance and reaction speed. Efficiency was computed for each participant and sound category, and statistical significance was assessed with a rmANOVA using participant group as a between-subject factor and stimulus roughness as a within-subject factor.

### Results

Average hit rate across participants and condition was H=.78 (SD=.18) and response time was RT=1.04s (SD=0.94). There was a main effect of stimulus roughness on efficiency, where the spatial location of rough-manipulated sounds was detected more efficiently than that of original sounds (Figure 3; mean efficiency difference M=0.31, 95% CI [0.14, 0.49], *F*(1,32)=6.65, *p*=.014, ges = 0.036). This difference was actually driven by accuracy: rough sounds were detected more accurately than original sounds (dprime: *F*(1,32)=6.15,*p*=.018), with no reduction of reaction time (z-score RTs: *F*(1,32)=1.10,*p*=.30). Importantly, the facilitating effect of roughness did not interact statistically with participant group (*F*(1,32)=0.52, p=.54), although paired t-tests only showed an effect of stimulus roughness in metal fans (mean efficiency difference M=0.40, 95% CI [0.05, 0.76], *t*(16)=2.46, *p*=.025), but not in non-fans (M=0.22, 95% CI [−0.13, 0.57], *t*(16)=1.24, *p*=.23).

**Figure 3.**
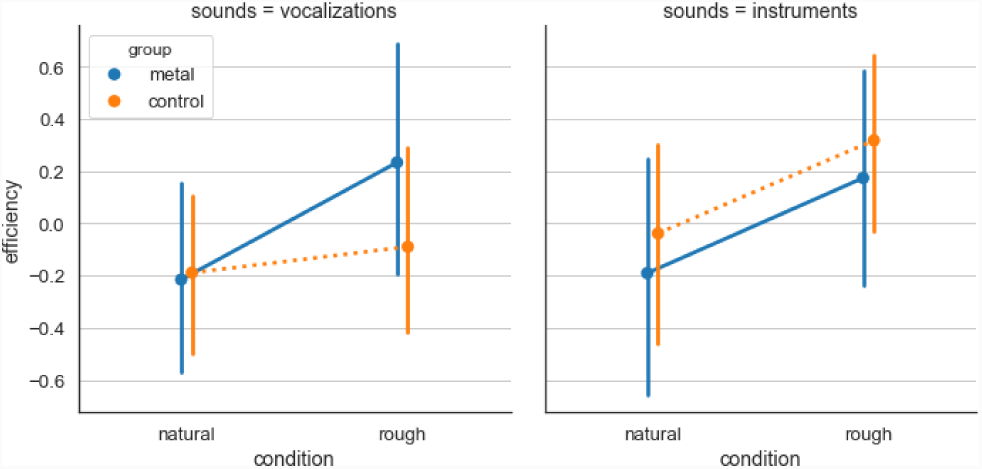
Effect of stimulus roughness on spatial localization (Exp.2), left: human vocalizations, right: musical instruments. Rough sounds were localized with more efficiency (more accurately at similar reaction times), and metal fans were no less sensitive to the facilitating effect of roughness than non-metal fans.

As an additional non-registered analysis, we also examined the effect of sound category (vocalization or instrument) on the efficiency of spatial localization: regardless or roughness, musical instruments were detected more accurately (mean difference of z-score d-prime: M=0.74, 95% CI [0.48, 1.01], *F*(1,32)=17.01, *p*=.0002, ges = 0.19), but also more slowly as compared to vocalizations (mean difference of z-score RTs: M=0.50, 95% CI [0.41, 0.60], *F*(1,32)=59.9, *p*=8.10^*-*^9, ges = 0.63), with the result of no effect on combined efficiency (Figure 3; *F*(1,32)=0.38, *p*=.54). None of these effects interacted with roughness, nor with participant group.

### Discussion

Our data replicates the previous finding that cues of vocal arousal facilitate the spatial localization of both vocal and musical sounds. Arnal et al. (2015) found that rough sounds were detected with both better accuracy and faster response time; on a similar task, our participants gave here more accurate responses with similar response times than for control sounds. It is possible that the latency effect additionally found by Arnal et al. (2015) is due to their making the baseline task more difficult by embedding target sounds in white noise at 5dB SNR, and adding a sinusoidal ramp of amplitude in the initial 100ms of the sounds. It is therefore significant that, even in ecological listening conditions, cues of roughness improve the accuracy of spatial localization.

Critically for our hypothesis, however, metal fans did not behave with less efficiency than non-metal fans when localizing rough sounds; if anything, they were even more accurate than non-fans. Consistently with Experiment 1, this pattern of result does not support the idea that extreme music lovers do so because they do not respond as intensely to cues of auditory threat: explicitly, they rate them as similarly negative and, implicitly, react to them equally fast and accurately as non-fans.

Olsen et al. (2018) found that word recognition accuracy for death metal lyrics was significantly enhanced for metal fans (65%) relative to non-fans (51%), which they interpreted as listener expertise in the acoustic features of metal vocalizations and their characteristic growl-like timbre. In a similar fashion, this expertise may here serve to amplify the sensory inputs to threat response circuits, and provide fans with a strong encoding of the features of auditory roughness in the subcortical auditory pathway (see e.g. Strait et al. 2012).

## Experiment 3: loaded preference task

Results from Experiment 1 and 2 do not give empirical support for a differential functioning of low-level threat response circuits in metal fans, who react equally negatively (Experiment 1), equally fast and even more accurately (Experiment 2) to cues of auditory threat in vocal and instrumental contexts than non-fans. Whether autonomic/behavioural threat responses and subjective fear are the result of two entirely orthogonal systems (LeDoux and Pine 2016) or the result of a unique fear generator with distinct effectors that can be independently modulated (Fanselow and Pennington 2018), it therefore appears that metal fans do not respond to cues of auditory threat in extreme music with an unusual variant of a fixed subcortically-determined behavior. As proposed above, an alternative hypothesis is that higher-order, top-down modulation by prefrontal cortical systems plays an important role in the aesthetic musical experience (Belin and Zatorre 2015).

A wealth of behavioural and neural data documents top-down contributions of executive functions and prefrontal systems to the prepotent processing of affective stimuli (Van Dillen et al. 2009; Greene et al. 2008; Abitbol et al. 2015), and show that these functions can be experimentally manipulated with dual-task paradigms. For instance, Gilbert et al. (1993) uses a visual digit-search task in which participants are instructed to press a response key each time the digit ‘5’ appears in a stream of rapidly-scrolling digits, while they concurrently read crime reports that contain both true and false statements; participants under such cognitive load were more likely to misremember false statements as true. Similarly, Greene et al. (2008) found that performing a concurrent digit-search task selectively interfered with utilitarian moral judgment (approving of harmful actions that maximize good consequences) but preserved non-utilitarian judgements based on emotional reactions (disapproving of harmful actions, regardless of outcome). Here, we use a dual-task paradigm in which participants listen and rate their preference for both metal and pop music extracts while engaging concurrently in a demanding digit-search task. With this paradigm, we test whether metal-fans’ positive orientation towards violent music is the result of cognitive control over inputs from more automatic first-order circuits which, as seen in Experiments 1 and 2, would otherwise predict the same negative reactions as in non-metal fans.

### Methods

#### Participants

Experiment 3 included the same N=34 participants (metal: 17: non: 17) as Experiments 1 and 2.

#### Stimuli

Stimuli consisted in 80 short (7-9s.) extracts from commercial musical songs of the metal (40) and pop music (40) genres. Songs in both genres were selected on the basis of participant responses to the screening questionnaire (see Experiment 1), using the following procedure: each participant of the metal (resp., pop) group listed 3 favorite titles of that genre; a list of 20 titles was selected from all of the participants’ responses with the criteria to include music that had (resp., did not have) clear cues of auditory threat (growl-like vocals, distorted guitars, noise and non-linearities) ; each title was then substituted by another similar, but lesser known song of a different artist using the “song radio” tool of the commercial music service Spotify*. The popularity of a given title or artist was estimated using Spotify’s “play count” for that title or that artist (for a similar methodology, see e.g. Bellogin et al. 2013). Substitute titles were selected if their play count was less than 10% of that of the most popular title of the most popular artist of the genre, and if their artist’s play count was less than 10% of that of the most popular artist of the genre. The rationale for the procedure was to select songs that were maximally similar to the group’s self-reported favorite items, but unlikely to be known/recognized by the participants. Finally, two 7- 9s. extracts from each of the 20 songs was selected, to be presented in each of the two experimental blocks (load/no-load), so that stimuli were matched in terms of musical content but not exactly repeated. The procedure resulted in 80 extracts (2 extracts × 20 songs × 2 genres), the same for all participants. Song list available in supplemental material S1.

#### Procedure

The experimental procedure consisted in two blocks of 20 trials, with and without cognitive load (counterbalanced across participants). In each block, trials consisted in pairs of musical stimuli (one of each genre), presented in a random order with a 1.5 s. inter-stimulus interval. Participants listened to the stimuli over headphones (Beyerdynamics DT770). Upon hearing the second stimulus of each pair, participants were instructed to report their preference for one or the other extract (2AFC), as well as a measure of their confidence in that preference (from 1 - “not at all confident” to 4 “very confident”).

In the load condition, streams of colored (red, green, blue, yellow) digits scrolled on the screen during each trial. The stream started 3s. before the beginning of the first musical excerpt, and continued until participants were prompted for a confidence rating. This ensured that both listening and music preference were done under the concurrent task, while confidence judgments were provided without cognitive load. Participant were instructed to press a key when digit 5 was presented on the screen in either red, green or yellow, but to inhibit their response if it was presented in blue. Digit probability was set at 0.3 for digit 5, and 0.1 for digits 1-4,6- 8; color probability was 0.4 for blue, and 0.2 for red, green and yellow. Digits were displayed at a fixed period in the range [200,300]ms, calibrated for each participant using an adaptive procedure (see below). To increase task demands, a warning message was displayed at each detection error (miss or false alarm).

In the non-load block, the same string of digits was presented on the screen, but participants were instructed to simply ignore them and focus on the main task. The order of the blocks was counterbalanced across participants, and the stimuli were pseudo-randomly assigned to one block or the other so that excerpts of the same songs appeared in different blocks.

The calibration procedure for digit search frequency was a 2-up-1-down staircase, aiming for a 70% detection rate. The initial period was set at 500ms, and the step size at 50ms. Throughout the procedure, period values were constrained to a minimum of 200ms and a maximum of 300ms. The procedure stopped after 12 inversions, and the final period was computed as the average period over the last 5 steps.

#### Preregistered analysis strategy

Participants’ preferences over the 20 trials of each block were aggregated into a score of preference for metal, by dividing the number of metal songs preferred over their alternative pop songs by the total number of trials (20). We then tested the effect of participant group (between-participant, 2 levels: metal/control) and condition (within-participant, 2 levels: load/control) on preference for metal and confidence scores using a rmANOVA.

### Results

Predictably, there was a large main effect of participant group on preference, with metal fans expressing stronger preference for metal over pop music alternatives independently of cognitive load ((Figure 4-top; mean difference of preference M=0.50, 95% CI [0.42, 0.57], *F*(1,32)=84.19, *p*= 1.78.10^-10^, ges=0.67). There was a main effect of cognitive load on participant’s response times and confidence, with slower (mean increase of RT M=480ms, 95% CI [280, 670], *F*(1,32)=11.5, *p*=.0019, ges=0.09) and less confident (mean loss of confidence M=−0.18pt on a 1-4 scale, 95% CI [−0.26, −0.09], *F*(1,32)=9.32, *p*=.004, ges=0.03) responses made under load, suggesting that our experimental manipulation indeed loaded cognitive functions. However, there was no main effect of the cognitive load manipulation on metal preference (Figure 4-top; mean loss of preference M=−0.01, 95% CI [−0.05, 0.03], *F*(1,32)=0.12,*p*=.72) and, critically for our hypothesis, no significant interaction between group and cognitive load (*F*(1,32)=2.92, *p*=.09).

**Figure 4.**
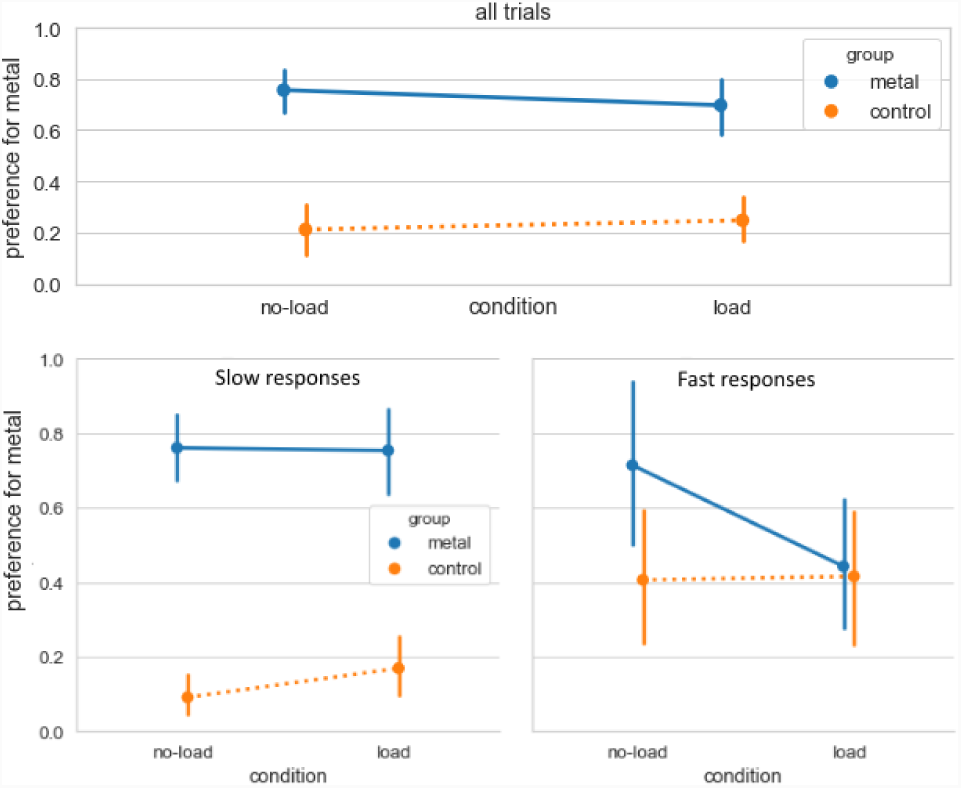
Effect of cognitive load on preference for metal music, in both metal fans and non-fans (Exp.3), top: all trials, bottom left: trials with slow responses, bottom right: trials with fast responses. While cognitive load had no effect on slow responses, the manipulation had an effect on preference responses when they were reported before the end of the second song (’fast responses’), with metal-fans reporting 36% less preference for metal over pop music while under load, while pop-fans did not show such a change in their musical preferences across the two conditions.

While our preregistered strategy for analysis failed to reveal any effect of cognitive load on participant preference, an additional exploratory analysis showed that participant preference response times were in fact bimodally distributed, with 25.8% of “fast” responses made while listening to the second song in a trial (before it was completely heard, i.e. <300ms post-song), and 74.2% of “slow” responses made after both songs were completely heard (i.e. >300ms post-song). We therefore grouped preference scores in fast/slow response types, and found that, while no effect of the cognitive load was observed in slow responses, the effect that we predicted initially was present in fast responses (Figure 4- bottom). For these trials, cognitive load reduced preference for metal in metal fans by 36% (mean loss of preference M=− 0.36, 95% CI [−0.61, −0.11], *t*(12)=−3.18, *p*=.008), while it did not affect preference for pop music in the control group (mean change of preference M=0.08, 95% CI [−0.18, 0.35], *t*(22)=0.66, *p*=.51)^†^.

### Discussion

Our dual-task paradigm with a taxing visual digit-search task was successful in creating cognitive load, as evidenced by 480ms-slower and less confident reports of musical preference in the concurrent music listening task. This pattern of result is weaker but consistent with previous paradigms of the same kind: with a slightly faster rate of digit display (140ms) but a simpler task (without inhibiting targets of certain colors) and a different domain of evaluation (moral choices), Greene et al. (2008) report a 750ms increase of response time; in Lee et al. (2007), a concurrent auditory task created a loss of confidence in visual judgments, with an effect size (d=0.5) also greater than what we find here.

Our data provides mixed evidence for the role of cognitive load in evaluating preference for metal music. While we found no effect of cognitive load on participants’ preference judgments for extracts of the metal or pop music genre in our preregistered analysis strategy, we found an effect of load when we restricted the analysis to those trials which participants answered rapidly (before the two extracts of a pair were played integrally). While this concerns only 25% of the data, that cognitive load impacted only fast responses is not incompatible with the literature. In van Dillen and van Steenbergen (2018), participants were time-limited and pressed to respond quickly to loaded trials (pictures of edible vs non-edible food) to avoid that participants engage in avoidant gaze strategies that could reduce interference with the digit-span task; in van der Wal and van Dillen (2013), they were instructed to drink liquid samples *all at once* before evaluating them. That cognitive load did not interfere with slower, self-paced responses may indicate that our visual cognitive-load task only had a relatively moderate impact on executive functions, and that slow trials correspond to those in which the cognitive load was only partial and did not prevent our participants from engaging higher order cognition during their judgement (Lavie 2010). It is also possible that load interfered as expected with sensory processing during listening, but that additional time taken after the direct experience of the stimuli allowed participants to engage in additional cognitive processes, such as semantic or autobiographic memory (e.g., “this is metal, and I like metal”), that may not have been impacted by our cognitive-load task. Further work should attempt to replicate this pattern of data with a paradigm involving higher cognitive load, and/or speeded responses of music preferences.

While the fact that cognitive load reduced a proportion of music preference towards metal in metal fans seems to indicate a role of controlled cognitive processes in preference for this musical genre, an alternative explanation is that load simply made participants unable to do the task : while speeded preference for metal music in metal fans was degraded under load to 0.42 (i.e. they on averaged preferred pop to metal), this proportion did significantly differed from the 0.5 chance level. However, this alternative interpretation is not really compatible with the fact that load did not degrade preference for pop music in the control group. Another possibility is that it was speeded judgments, rather than load, which ‘regressed’ preferences toward the mean, but this interpretation is also made unlikely by the fact that, even in theses responses, metal fans had marked preference for metal in the no-load condition.

## General discussion: towards a higher-order theory of the emotional experience of music

While it is generally admitted that the cognition of musical signals is continuous with that of generic auditory signals (Schlenker 2017) and that, in particular, the emotional appraisal of music largely builds on innately-programmed, primary-subcortical brain systems evolved to respond to animal signaling (Blumstein et al. 2012), human prosody (Juslin and Laukka 2003) and environmental cues (Ma and Thompson 2015), the case of appreciation for extreme metal music seems a theoretical conundrum (Thompson et al. 2018). It could be that metal fans differ from non-fans in how they process threat signals at the subcortical level, showing deactivated or reconditioned responses that differ from controls - a view that has lead some to call appreciation for violent music a psycho-social dysfunction (Stack et al. 1994; Bodner and Bensimon 2015; Sun et al. 2017). However, from a more recent higher-order perspective of emotional experience (LeDoux and Brown 2017), it is also possible that fans’ appreciation for metal reflects the modulation/inhibition by the cortical circuits of higher-order cognition of an otherwise normal low-level response to auditory threat. In a series of three experiments, we have shown here that, at the perceptual and affective levels, metal fans react in fact equally negatively (Experiment 1), equally fast and perhaps even more accurately (Experiment 2) to cues of auditory threat in vocal and instrumental contexts than non-fans, and that, under some conditions of speeded preference judgments, cognitive load reduces fans’ appreciation of metal to the level experienced by non-fans (Experiment 3). Taken together, these results provide no support to the idea that extreme music lovers do so because of a different low-level response to threat, but rather highlight a critical contribution of higher-order, controlled cognitive processes in their aesthetic experience.

While these results have implications for a growing corpus of psychological studies of metal music (Gowensmith and Bloom 1997; Bodner and Bensimon 2015; Sun et al. 2017; Thompson et al. 2018; Olsen et al. 2018), notably confirming that viewing metal as dysfunctional “problem music” is empirically untenable, implications for the general theory of musical emotions are, in our view, even greater. They shape a model of musical emotions which significantly extends the traditional view, in which the cortical and subcortical signals sent by affective and sensory systems (auditory thalami, auditory cortices) do not simply feed forward relatively unaltered to associative cortices (following e.g. right temporal-frontal pathway of emotional prosody processing-Schirmer and Kotz 2006), but can also be thoroughly modified/inhibited by the circuits of higher-order cognition, to the point of creating emotional experiences (e.g. here, liking the music; in Thompson et al. 2018, the experience of peace or joy) that appear to contradict the low-level cues that serve as input to these evaluations (e.g. here, auditory roughness). What is significant in the present pattern of results is that behavioural signatures of both types of responses simultaneously co-exist in the system: metal fans exhibit both ‘typical’ low level processes that appraise rough sounds as negative and worthy of immediate attention (Experiments 1 and 2) *as well as* high-order systems able to assert cognitive control over these responses and produce positive emotional experiences (Experiments 3).

This model suggests that there is in fact a hierarchy of emotional experiences to music. Some, like that of rejecting metal music as threatening and violent, are strongly conditioned by low-level systems and flow relatively unaltered into conscious awareness. Others, like appreciating metal, are significantly reshaped by cognitive control and culturally situated learning. It is perhaps ironical that positive responses to metal, once dismissed as dysfunctional or unsophisticated, may be one of the most cognitively refined in this spectrum of experiences. Other reactions of the same nature may include positive reactions to sad music (Vuoskoski et al. 2012), or negative emotions to entraining, happy music (e.g., *“Even if some culturally-determined part of your mind is saying ‘I hate this song’, your body will ecstatically sing along with Debby Boone in ‘You light up my life’* - Oswald 2000).

This idea that low-level responses shaped by evolution, and higher-order responses shaped by the social environment coexist and interact provides a unified framework to think about the interactions between biological and cultural evolution in the shaping of modern human musical experience (Bryant 2013): sound patterns such as the distorted guitar sounds and harsh vocals of metal music exploit evolved perceptual response biases manifested in first-order systems, but then take on distinct/controlled emotional values through cultural evolutionary processes, reflected in higher-order responses. This model also brings musical emotions in line with modern constructionist views of emotions (Barrett 2017; Cespedes-Guevara and Eerola 2018), for which the emotional experience is a psychological event constructed from more basic ‘core affect’ and higher-level conceptual knowledge. For fans of violent or sad music, the psychological construction of a positive experience from negatively-valenced sensory cues may be similar to that of constructing “invigorating fear” from a roller-coaster ride or “peaceful sadness” from enjoying a moment of solitude after a busy day (Wilson-Mendenhall et al. 2013).

More importantly, several predictions can be made from this model. First, because they implicate additional cognitive resources and less direct sensory evidence, one might expect that higher-order musical experiences e.g. preference for metal or sad music should be both slower and less confident than lower-order musical experiences e.g. dislike for metal or preference for pop music. In our data (Experiment 3), although this may reflect population rather than meaningful differences, judgments of preference for metal in metal fans were non-significantly slower (M=−239ms, 95% CI [- 618ms, +139ms], t(32)=1.28, p=.20) but significantly less confident (M=−0.37, 95% CI [−0.69, −0.05], t(32)=−2.37, p=.02) than judgments of preference for pop in non-fans. Further work should examine these differences in a within-subject, one-interval task more appropriate to measuring reaction times. Second, because first- and higher-order responses are assumed to co-exist during the emotional experience, one would expect to measure physiological reactions (e.g. pupil dilation Oliva and Anikin 2018) or neural activity (in, e.g., the amygdala Arnal et al. 2015) indexing normal response to threat relatively independently of the listener’s positive or negative emotional evaluation of metal music. Third, because executive functions involved in cognitive control are implemented in frontal lobe regions (Duncan and Owen 2000), one would expect that positive higher-order emotional reactions to e.g. violent or sad music should be degraded to more direct aversive responses with experimental manipulations such as transcranial magnetic stimulation to the dorsolateral prefrontal cortex (Tassy et al. 2011), or during sleep. Finally, at the population level, appreciation for metal, because it implicates controlled cognitive processes and executive functions, may be correlated with greater capacity for emotional regulation, just like appreciation for sad music may be correlated with greater trait empathy Vuoskoski et al. (2012).

Finally, while data from Experiment 3 seem to implicate controlled processes in the appreciation of metal, and less so in pop music, our results leave open many possibilities concerning the nature or timing of these processes. First, they leave the notion of ‘cognitive load’ relatively under-specified. Our task, a speeded digit search, loads both executive functions involved in updating (attention to novel digits) and inhibiting (inhibiting responses to targets of one specific color), but not e.g. in task switching (Miyake et al. 2000), and it is unclear which of these processes specifically contributes to the construction of the emotional experience. Second, the present results do not address the appraisal mechanisms that govern the emotional responses that, according to our theory, support liking metal. Processes inhibited in Experiment 3 could involve, e.g., focusing one’s attention on other features of the music than threatening cues (e.g. treating growling vocals as a non-emotionally-significant singing style, and focusing instead on on words or melody - Olsen et al. 2018), engaging in psychological distancing (e.g. evaluating metal sounds as a virtual threat that presents no actual danger to personal safety - Menninghaus et al. 2017), establishing an aesthetic judgmental attitude (Brattico and Vuust 2017), or recontextualizing cues of violence as not directed toward the self, but from the self toward an hypothetical other (explaining e.g. feelings of power often reported by metal fans - Thompson et al. 2018). Finally, here we took music preference as a proxy for emotional experience, but preference is mediated by many variables other than a positive affective response, including imaginal and analytical responses (Lacher and Mizerski 1994), which all could have been affected by our load manipulation. Further work should therefore attempt to replicate the effect of cognitive load on more direct and varied measures of emotional experience.

## Supplementary material

### S1: Song extracts used as stimuli in Experiment 3

#### Metal group

Deez Nuts - *Purgatory*. ©2017 Century Media Records

Enterprise Earth - *Shroud of Flesh*. ©2017 Stay Sick Recordings

Veil of Maya - *Fracture*. ©2017 Sumerian Records

Lordi - *How to Slice a Whore*. ©2014 AFM Records

Testament - *The Pale King*. ©2016 Nuclear Blast

The Color Morale - *When One was Desolate*. ©2009 Rise Records

Heaven & Hell - *I - Live*.©2007 Rhino Entertainment

Erra - *Skyline*.©2016 Sumerian Records

Carcass - *Edge of Darkness*.©1996 Earache Records

Soil - *Way Gone*.©2017 Pavement Entertainment

Coal Chamber - *Entwined*.©1999 Woah Dad!

Between The Buried and Me - *The Coma Machine*.©2015 Metal Blade Records

Testament - *Trails of Tears*.©1994 Atlantic

Coal Chamber - *Beckoned*.©2002 Woah Dad!

Deathstars - *Death Is Wasted On the Dead*.©2014 Deathstars

Death - *Spirit Crusher*.©2011 Relapse Records

Miss May I - *Never Let Me Stay*.©2017 Sharptone

Nonpoint - *Be Enough*.©2016 Spinefarm Records

Allegaeon - *From Nothing*.©2016 Metal Blade Records

Miss May I - *Crawl*.©2017 Sharptone

#### Control group

Zaho - *Te amo*. ©2013 Parlophone Records

Lea Michele - *Empty Handed* ©2013 Columbia Records

Loreen - *Statements*. ©2017 Warner Music

Benjamin Ingrosso - *Dance You Off*. ©2017 Record Company TEN

S Club 7 - *I’ll Keep Waiting*. ©2000 Polydor Ltd

Hilary Duff - *Rebel Hearts*. ©2005 Hollywood Records

Amerie - *Hatin’ On You*. ©2002 Sony Music Entertainment

Vrit - *Solutions*. ©2017 Vrit

Elder Island - *Key One*. ©2016 Elder Island

Superbus - *On the River*. ©2012 Polydor

S Club 7 - *Dance Dance Dance*. ©2001 Polydor Ltd

Tt - *Chanteur Sous Vide*. ©2016 Fffartworks

Tony ! Toni ! Ton ! - *My Ex-Girlfriend*. ©1993 PolyGram

Athlete - *Airport Disco*. ©2016 Chrysalis Records

Thirteen Senses - *Thru The Glass*. ©2004 Mercury Records

Hollysiz - *Rather Than Talking*. ©2017 Hamburger Records

vrit - *Somewhere in Between*. ©2017 Vrit

Rupaul - *Kitty Girl*. ©2017 RuCo

Rupaul & Ellis Miah - *Just a Lil In & Out*. ©2017 RuCo

### S2: Preregistration document

Document submitted to the Ecole Normale Supérieure (ENS) Cogmaster office, Decision dated Feb 1st, 2018.

## Acknowledgements

All data collected at the Centre Multidisciplinaire des Sciences Comportementales Sorbonne Universit - INSEAD. Author contributions: LO, LG, JJA designed the experiment. ML developped the ANGUS stimulus manipulation. LO and LG collected the data. LO, LG and JJA analysed the data. JJA wrote the manuscript, with contributions from LG.

## Footnotes

* https://www.spotify.com, data accessed: March 2018

† A statistical note: analyses in the slow and fast response subgroups were done with independent rather than paired t-tests between the load and no-load condition (despite some amount of shared variance within some of the participants), because not all participants had slow and fast responses in both load conditions. If restricting the analysis to those participants who had fast responses in both load conditions, a repeated-measure ANOVA showed a similar group × load interaction (*F*(1,11)=7.01,*p*=.022), and a similar reduction of preference for metal in the metal group (mean loss of preference M=−0.40, 95% CI [−0.70,−0.09]), but that group included only n=4 metal fans.

